# Self-organised segregation of bacterial chromosomal origins

**DOI:** 10.1101/304600

**Authors:** Andreas Hofmann, Jarno Mäkelä, David Sherratt, Dieter Heermann, Seán M. Murray

## Abstract

In spite of much effort, many aspects of chromosome organisation and segregation in bacteria remain unclear. Even for *Escherichia coli*, the most widely studied bacterial model organism, we still do not know the underlying mechanisms. Like many other bacteria, the chromosomal origin of replication in *E. coli* is dynamically positioned throughout the cell cycle. Initially maintained at mid-cell, where replication occurs, origins are subsequently partitioned to opposite quarter positions. The Structural Maintenance of Chromosomes (SMC) complex, MukBEF, which is required for correct chromosome compaction and organisation, has been implicated in this behaviour but the mode of action is unknown. Here, we build on a recent self-organising model for the positioning of *E. coli* MukBEF, to propose an explanation for the positioning and partitioning of origins. We propose that a specific association of MukBEF with the origin region, results in a non-trivial feedback between the self-organising MukBEF gradient and the origins, leading to accurate positioning and partitioning as an emergent property. We compare the model to quantitative experimental data of origin dynamics and their colocalisation with MukBEF clusters and find excellent agreement. Overall, the model suggests that MukBEF and origins act together as a self-organising system for chromosome segregation and introduces protein self-organisation as an important consideration for future studies of chromosome dynamics.

## Introduction

The faithful and timely segregation of genetic material is essential for all cellular life. In eukaryotes the responsibility for chromosome segregation lies with a well-understood macromolecular machine, the mitotic spindle. In contrast, the mechanisms underlying bacterial chromosome segregation are much less understood but are just as critical for cellular proliferation. Nevertheless, a lot has been learned in recent years (see [1] for a review). In particular, the starting point for (bidirectional) chromosomal replication, the origin, has been found to have a crucial role in chromosome organisation and segregation. Not only is it duplicated and segregated first but its genomic position defines other macrodomains [2] and its spatial position may determine the overall organisation of the chromosome [3].

The positions of genomic loci, including the origin, change as replication and segregation progress. Initiated at the chromosomal origin, replication proceeds bi-directionally down each arm of the chromosome. In slowly to moderately growing (1-2 hr generation time) *Escherichia coli* cells, the home positon of the origin, *oriC* (henceforth and in the model, *ori*), in new-born cells is at mid-cell [4–6]. After 10-15 min of ‘cohesion’, in at least part arising from interlinking of the two daughter chromosomes (precatenation) [7–12], duplicated origins separate and migrate, initially very rapidly, to opposite quarter positions, which become the new home positions for the remainder of the cell cycle [13,14]. Other genomic loci migrate sequentially to independently acting replisomes [9,11,15,16] and subsequently segregate with similar dynamics [14]. However, in spite of extensive research, the forces underlying this behaviour are unknown.

Many bacterial species carry ParABS (Par) systems on their chromosomes [17]. First discovered in low-copy number plasmids, where they are required for correct partitioning, they also play a role in chromosome segregation. However, they are typically dispensable during normal growth [18–21] (*Caulobacter crescentus* is a notable exception [22]). Moreover, even in these cases, the chromosomal origins still show non-random positioning albeit perturbed or altered [20,23,24]. Therefore, it is clear that there are alternative mechanisms for origin positioning yet to be discovered. This is especially the case for bacteria that do not carry chromosomal Par systems such as *E. coli*, the most widely studied prokaryotic organism, and many of its relatives within the gammaproteobacteria.

Clues can likely be found in a group of DNA maintenance proteins that co-evolved within the gammaproteobacteria, including Dam, SeqA, MukBEF, MatP and MaoP [25,26]. In particular there is increasing evidence that MukBEF, a functional homolog of ubiquitous Structural Maintenance of Chromosomes (SMC) complexes [27,28], has a role in positioning origins in *E. coli*. Under slow growth conditions, MukBEF forms a small number of dynamic clusters (henceforth foci) located, in close association with *ori* at the middle or quarter positions [29,30] and the splitting and movement of these foci occurs concurrently with the segregation of *ori* to the quarter positions [10,29]. Depletion of MukBEF results in *ori* mis-positioning, which is restored upon repletion [30], suggesting a connection between their positioning. MukBEF recruits the type II topoisomerase Topo IV [10,31], which is required for the timely removal of catenanes from newly replicated sister chromosomes [11]. Further support that MukBEF foci positions *ori* rather than the other way around came from the discovery that depletion of Topo IV results in catenation and mis-positioning of *ori* but not of MukBEF foci with decatenated *ori*s re-associating with MukBEF foci after restoration of TopoIV activity [10].

However, up to recently the idea that MukBEF positions *ori* suffered from a fundamental setback. If MukBEF foci position *ori*, what positions MukBEF? Given that foci turnover continuously on a timescale shorter than a minute [32] and that MukBEF binds DNA non-specifically, how does it even form foci? We recently proposed a solution to this conundrum in the form of a self-organising and self-positioning mechanism based on phase-fixing a stochastic Turing pattern [33] (see Figure S1a and the methods section for a review). With this model in hand, we are now in a position to test the hypothesis that MukBEF positions chromosomal origins. In particular, we are asking whether the self-organising MukBEF gradient (as described in our previous model) has the correct properties to act as an attracting gradient for *ori*. Additionally, it is critical that *ori* are correctly partitioned, i.e. each *ori* is recruited to a *different* MukBEF focus, a non-trivial requirement.

We find that a self-organising MukBEF gradient can indeed accurately reproduce the observed *ori* dynamics, apparent diffusion constant and drift rate. The specific loading of MukBEF within *ori* introduces a non-trivial interaction between MukBEF foci and *ori* that leads to accurate and stable partitioning as an emergent property of the system. Importantly, the model does not contain any actual directed force. MukBEF requires energy in the form of ATP to establish a self-organised gradient but it is not pulled to the middle or quarter positions by any active force. Similarly, the attraction of *ori* up the MukBEF gradient is due to energetic considerations and the elastic nature of the chromosome (a DNA relay [34]) resulting on the macro scale in an effective (rectification) force and directed motion.

## Results

As discussed above, multiple lines of evidence point to an interaction between MukBEF foci and *ori* suggest that MukBEF foci position *ori*. How could this interaction be mediated? In *Bacillus subtilis*, SMC, which also colocalises with *ori* [35], is loaded onto the DNA at *parS* sites located mostly in the origin region by the protein ParB [35–37] (see [38] for a review of SMC loading, translocation and function). However, MukBEF and SMC do not show high specific DNA binding [12,36] likely due to their dynamic nature and ability to translocate along the DNA. Thus those studies relied on prior knowledge of ParB-*parS* (similarly for MukBEF unloading from the terminus region by MatP-*matS* [12]). It is therefore conceivable that a loading site/protein exists within the origin region of *E. coli* but has not been identified. We take the presence of such a specific loading site as the starting assumption of the model and we use ‘site’ without prejudicing the number of sites or the existence of a loading factor. Alternatively, and notwithstanding the lack of evidence for high specific binding, the association between MukBEF foci and *ori* could arise as a consequence of *ori* being a ‘stop’ region for MukBEF complexes undergoing transport along chromosomes (see discussion below).

The nature of the attraction of *ori* to its home position has been characterised by Kuwada *et al.* using time-lapse fluorescence microscopy [13]. They found that the mean *ori* velocity exhibits a linear relationship with the long-axis position, indicative of a diffusing particle experiencing a harmonic (quadratic) potential, or equivalently, a damped spring-like force. Any model of *ori* positioning must be able reproduce this behaviour. Our assumption of a higher affinity association between MukBEF and *ori*, implies that the MukBEF concentration profile would as a potential energy surface for *ori* that is minimised when *ori* are located at peaks in the MukBEF concentration *i.e. ori* would move up MukBEF peaks.

### Direction movement of *ori* can arise from spatially-dependent looping probabilities

From a microscopic point of view, we envisage the movement of *ori* as being due to the elastic nature of DNA [39] allowing it to locally sample the MukBEF concentration and thereby bias its direction of movement as in the Par-based DNA relay model of *ori* segregation in *C. crescentus* [34]. The distinction here is that the protein gradient is self-organised, and that the cargo does not destroy the gradient as it moves. However, two effects were not considered in that model. First, the *ori* was treated as inherently freely diffusing (in the absence of interactions with the ParA protein gradient) while there is no reason to believe that it does not exhibit the same elastic fluctuations about its home position as other loci. Secondly, as the *ori* migrates across the cell, it will experience an entropic counter force due to the polymeric nature of the chromosome.

We therefore wanted to test the DNA relay hypothesis as applied to this system. In particular, we wondered whether directed movement of *ori* positioning can arise due to the non-uniform MukBEF concentration profile and its promotion of long-range DNA interactions [40]. We modelled the chromosome as a self-avoiding ring polymer confined in a rectangular cuboid and used the dynamic loop model [41] to mimic the formation of DNA loops between *ori* and distant DNA sites mediated by MukBEF. MukBEF molecules were not directly simulated, but were incorporated via a spatially dependent looping probability along the long axis of the cuboid (nucleoid) representing the MukBEF concentration profile.

We found that the steady-state distribution of *ori* positions matched the applied Gaussian looping probability distribution (Fig. 1a). This was to be expected, given that formation of long contacts with the *ori* would inhibit its mobility. The important question is whether this distribution arises in a directed manner or as a time-average. When we examined the velocity of the *ori* as a function of long-axis position, we found that the *ori* indeed experiences a restoring velocity towards mid-cell i.e. directed movement (Fig. 1b). This demonstrates that a spatially varying probability of *ori* to form long-range contacts with other loci is able to break the symmetry of the thermally driven elastic fluctuations resulting in an effective force and biased movement. This supports the plausibility of our assumption that *ori* is attracted up peaks in the MukBEF concentration.

**Figure 1:**
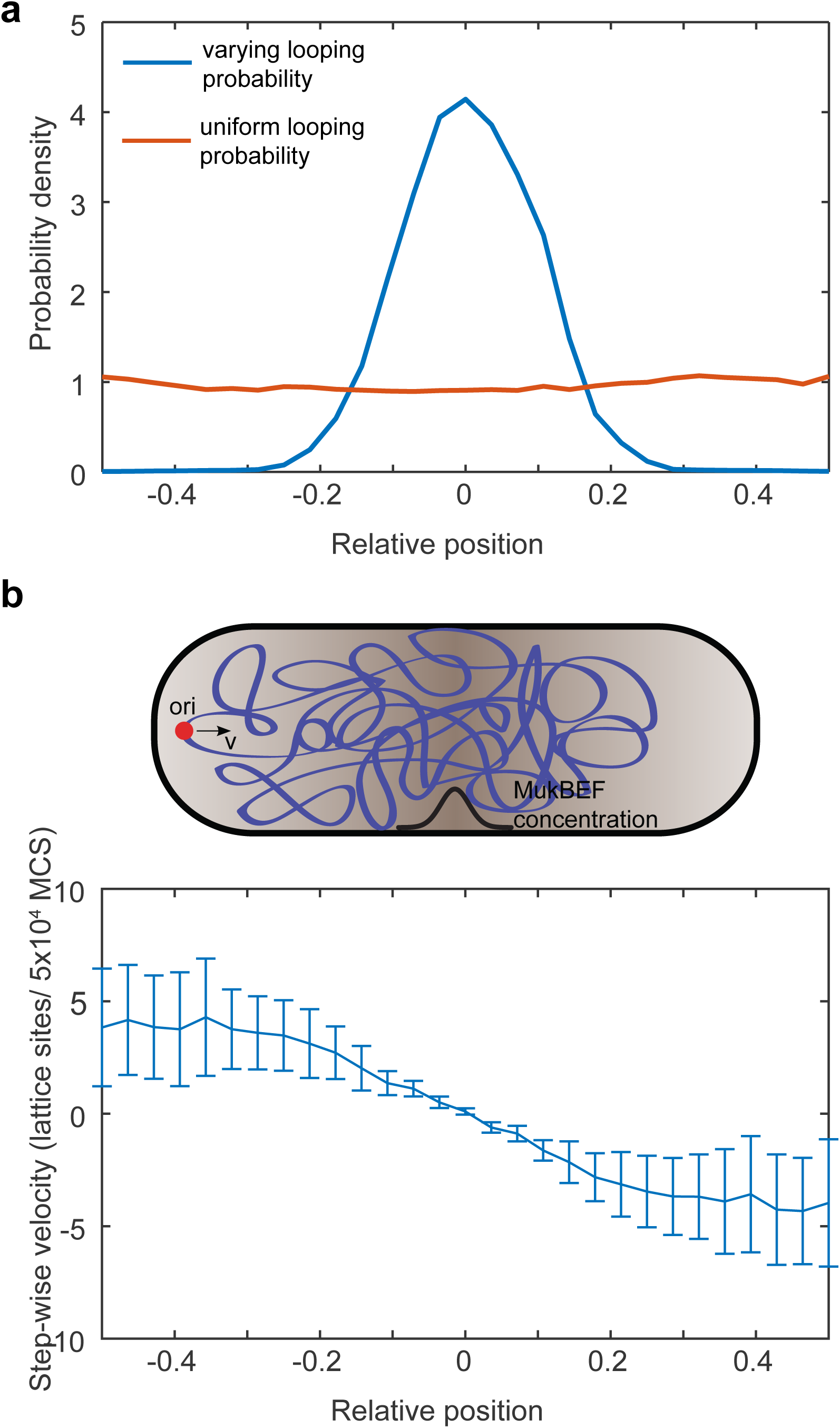
Directed movement of *ori* can arise from spatially dependent looping probabilities (a) Probability density of relative *ori* position along the long axis of the cuboid (aspect ratio 4:1) with (blue) and without (red) a spatially-varying looping probability (a Gaussian centred at 0 with standard deviation 0.1 in units of long-axis length. (b) (Bottom) The step-wise *ori* velocity along the long axis as a function of relative position along the long axis. Error bars indicate standard error. (Top) A diagram indicated the biased direction of movement. The *ori* position was read out every 50000 Monte Carlo time-steps (MCS). Data is from 10 independent simulations with approximately 10000 data points from each.

### Model of *ori* positioning by self-organised MukBEF reproduces mid-cell positioning

Given the self-organising and fluctuating nature of the MukBEF gradient in our model [33] (which is representative of the *in vivo* behaviour), it was not obvious that it would generate the observed linear *ori* velocity profile. However, we found that the steady-state distribution of MukBEF (we refer to the slowly-diffusing state that constitute foci [32,33] simply as MukBEF - see methods for details) closely resembles a Gaussian, i.e. the equilibrium distribution for a harmonic potential (Fig. S1b). Thus, MukBEF itself exhibits the same distribution as if it experiencing a damped spring-like force towards mid-cell. As a Gaussian very closely resembles a quadratic around its peak, MukBEF might therefore be able to act as an effective potential for *ori* and generate the linear velocity profile observed by Kuwada *et al* [13].

Based on the previous results, we therefore incorporated *ori* into our stochastic reaction-diffusion simulations by treating it is a diffusing particle, the movement of which is biased in direction of increasing MukBEF concentration (see methods for details). We took initially the case of short cells (2.5 µm) with a single *ori*. We found that *ori* track the self-organised MukBEF foci, resulting in mid-cell positioning (Fig. 2a,b and Fig. S1c). Furthermore, as suggested by the steady state MukBEF distribution, we found that simulated *ori* experience a linear restoring velocity (Fig 1c). A quantitative comparison of our simulations with the experimental data from Kuwada et al. [13] was carried out by fitting the data in the mid-cell region to a theoretical model of diffusion in a harmonic potential, thereby obtaining an apparent diffusion constant and drift rate. We used only cells with a single *ori* focus visible, which, unlike cells with two *ori*, can be grouped together without scaling by simply aligning them according to their mid-cell positions. We found that by adjusting the diffusion and drift input parameters, we were able to obtain excellent agreement with the experimental results (Fig 2c).

**Figure 2:**
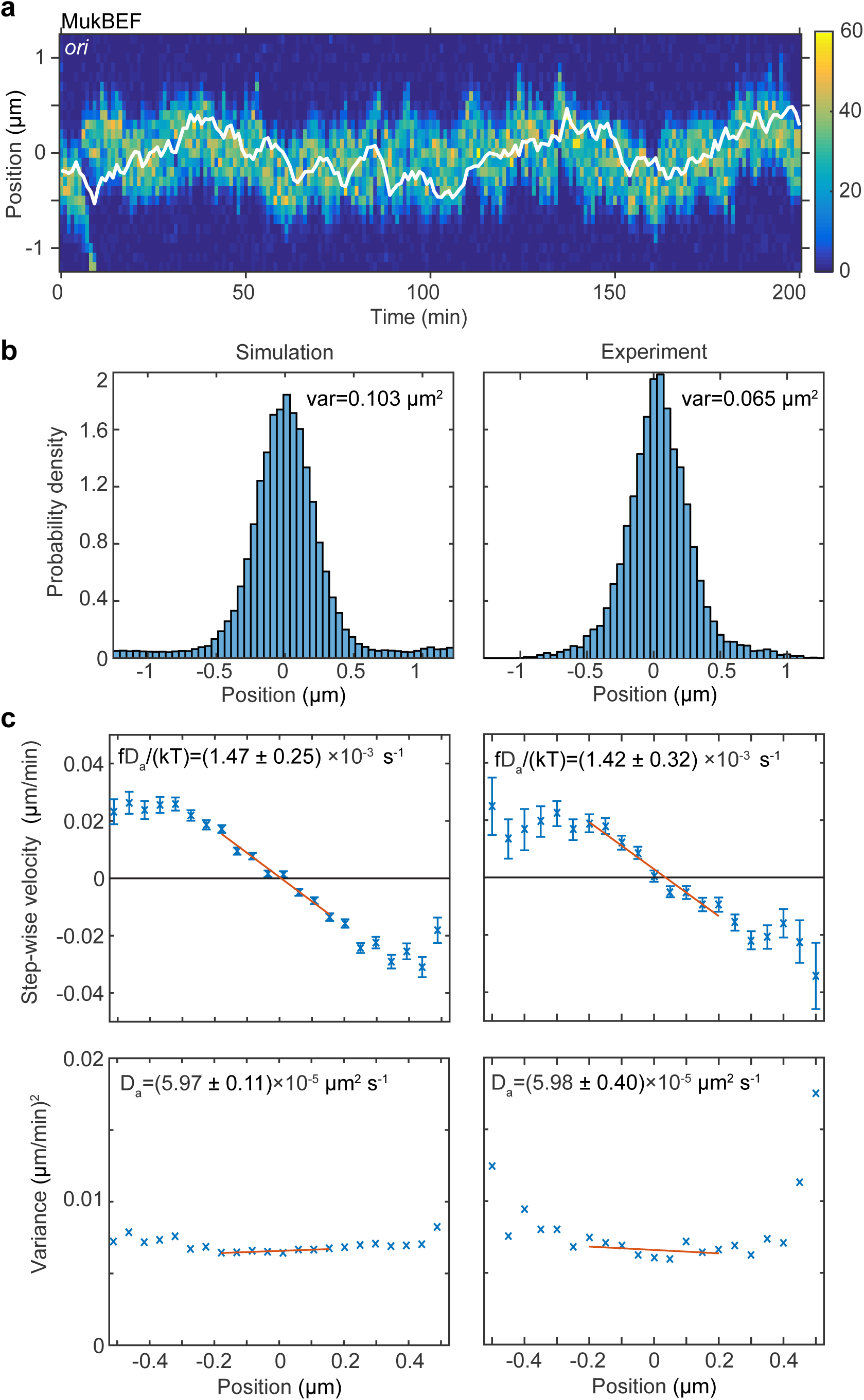
Ori positioning by a self-organised protein gradient reproduces experimental results (a) Kymograph from a single simulation showing the number of MukBEF molecules (colour scale) and the position of the *ori* (white line). (b) and (c) A comparison between the experimental data of Kuwada et al. [13] and the results of simulations in the case of a single *ori*. (b) Histograms of *ori* position (unscaled) along the long axis of the cell. Zero is the middle position. (c) Mean (top) and variance (bottom) of the step-wise velocity as a function of position relative to midcell. Bars indicate standard error. The linear velocity profile at mid-cell is indicative of diffusion in a harmonic potential 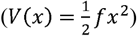. In such a model the variance of the step-wise velocity is independent of position. Thus we obtain the apparent diffusion constant, _a_ and drift rate 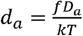 by fitting to the central region. Bounds are 95% confidence intervals. Red lines are weighted linear fits. Simulated data are from 100 independent runs, each of 600 min duration. Experimental data are based on over 16000 data points from 377 cells. Both data sets use 1 min time-intervals. Simulations are from 2.5um cells, whereas experimental data is from a range of cell lengths. See methods for further details and model parameters.

However, while promising, these results are not sufficient to suggest that MukBEF can explain the *in vivo* behaviour of *ori*. The challenge arises after the *ori* has been replicated. A true partitioning mechanism must ensure that each replicated *ori* moves to a *different* quarter position. A simple gradient based mechanism cannot, *a priori*, satisfy this requirement as both *ori* could just as easily move towards the same quarter position. Furthermore, the experimental data suggests that once *ori* separate they do not subsequently interchange their positions (cross paths). This ordering is essential during multi-fork replication, where the multiple *ori* of each segregated chromosome must be positioned to the appropriate cell half to avoid guillotining the chromosome upon cell division.

### Specific loading leads to stable and accurate partitioning

To examine if the model is capable of accurate and ordered partitioning, we performed simulations with two *ori* in longer cells of 5 µm, in which MukBEF self-organises into, on average, two foci, one at each quarter position. The average profile of *ori* positions displayed two peaks centred on the quarter positions. However, looking at individual simulation we found that in approximately half of the simulations both *ori* were near the same quarter position (Fig. 3a), clearly indicating that partitioning was not accurate.

**Figure 3:**
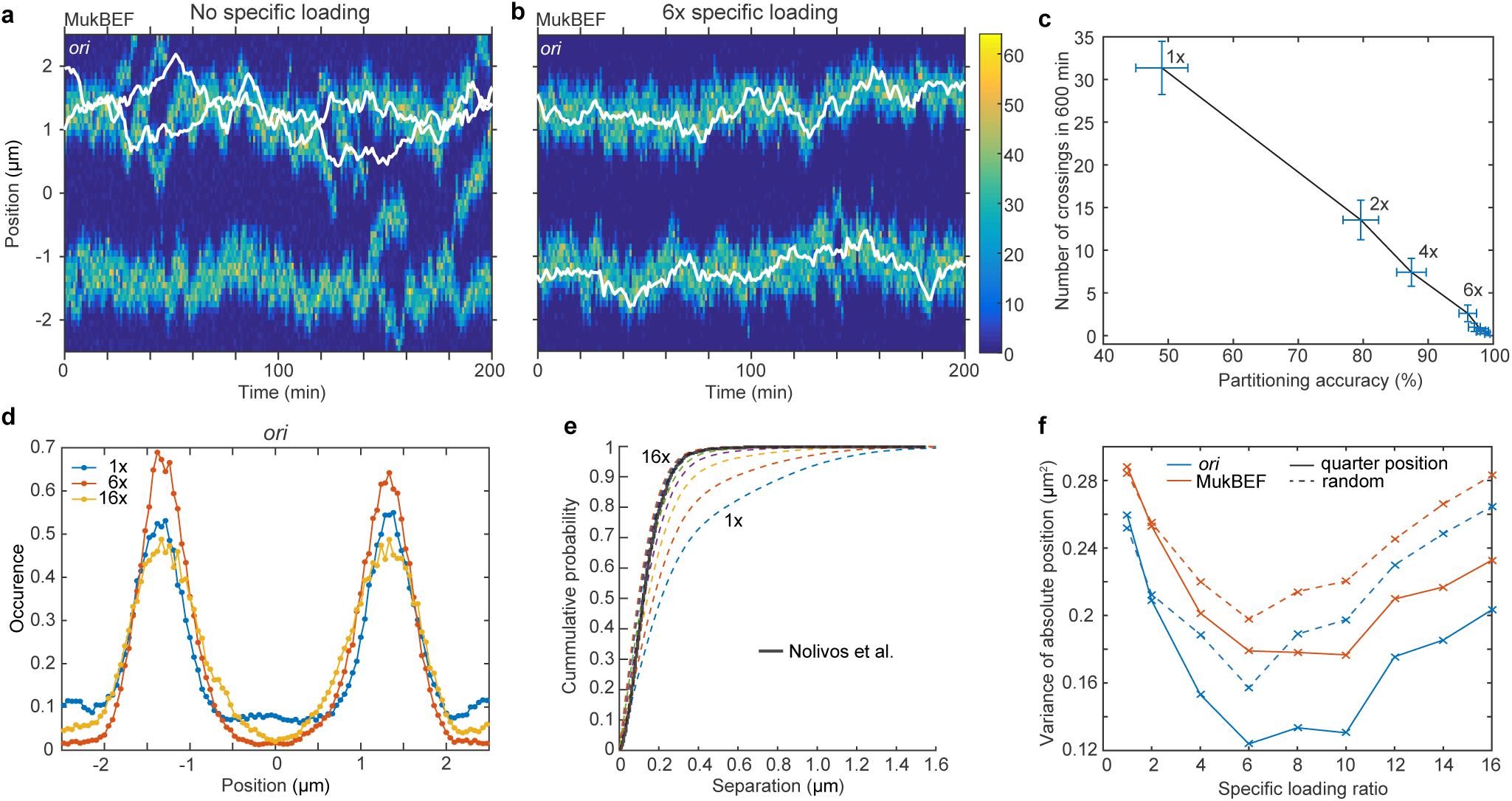
Specific loading of MukBEF at *ori* leads to correct and stable partitioning (a) Example simulated kymograph showing two *ori* (white lines) diffusing around the same MukBEF peak (colour scale). This occurs approximate 50% of the time. (b) The addition of 6x specific loading of MukBEF at *ori* positions results in correct partitioning of *ori*. The loading rate in each of the spatial compartments containing *ori* is six times that of the other 48 compartments. The total loading rate is unchanged. (c) Partitioning accuracy is measured by the fraction of simulations with *ori* in cell halves. Stability is measured by the number of times *ori* cross paths. Both partitioning accuracy and stability increase with specific loading up to approximately 6x. Specific loading ratios are as in (f). Points and bars indicate mean and standard error over independent simulations. See also Fig. S2c. (a) Histograms of *ori* positions for 1x, 6x and 16x specific loading. Positioning is more precise at 6x than with no or 16x specific loading. See Fig. S2a for MukBEF distributions. (e) The cumulative probability distribution for the separation distance between *ori* and MukBEF peaks. Experimental data (black line) is from Nolivos et al. [12]. The addition of specific loading leads to substantially better agreement. Specific loading ratios are as in (f). See also Fig. S1d. (f) The variances of individual peaks (obtained by reflecting the data around the mid-position) have a minimum at approximately 6x specific binding. Solid lines are from simulations with *ori* initially at the quarter positions, as for (a)-(e). Dashed lines are from simulations with random initial *ori* positions. Simulations were performed for a 5 µm domain and two *ori* but otherwise as in Fig. 2.

The previous simulation incorporated the movement of *ori* up the MukBEF gradient but *ori* had no effect on MukBEF. In previous work, we showed that a specific loading site at a fixed spatial location has an attractive effect on the positioning of self-organised MukBEF foci due to the modified flux differential across foci [33]. Thus the presence of a specific loading site in (or specific association to) the *ori* should lead to an effective mutual attraction between *ori* and MukBEF foci. We expected this to increase the association between the two but it was not clear what effect it would have on overall MukBEF-*ori* positioning. Surprisingly, we found that the addition of a specific loading at *ori* results in stable and accurate partitioning (Fig. 3b). A specific loading ratio greater than six (i.e. six times more binding than elsewhere) was sufficient to ensure that one and only one *ori* was positioned to each quarter position and they do not interchange (Fig. 3c,d). Furthermore, when we examined the colocalisation of *ori* with MukBEF foci as has been quantified *in vivo* [10,12], we found that similar levels of specific loading results in very similar cumulative probability distributions of the separation distance (Fig. 3e and S1d).

Additional effects were observed. The positions of both MukBEF and *ori* deviate less from the quarter positions and are almost never at mid-cell (Fig. 3d, Fig. S2a). However this effect was reversed at higher specific loading ratios and we found that six to ten times resulted in the narrowest distribution peaks (Fig. 3f). This did not affect partitioning accuracy or the number of times *ori* crossed paths which remained very high and small respectively (Fig. 3c). However, looking at individual simulations, we observed that the nature of the variance was different. At no or low specific loading, additional (more than two) transient MukBEF foci appear and disappear (Fig. 3a) resulting in a broader average distribution, while at high specific loading the number of foci is maintained accurately at two (Fig. S2b) and the foci are tightly associated to *ori* (Fig. 3e) but are more mobile than at intermediate ratios. We explain this effect and the mechanism behind accurate partitioning below.

Returning to shorter cells with a single *ori*, we measured the effect of specific loading on the apparent diffusion constant and drift rate and found that only the drift rate changes significantly (Fig. S3a,b). However, we are able to choose new values for the diffusion and drift parameters (at 6x specific loading) such that the agreement with experimental data was recovered (Fig. S3d). Furthermore, the position distribution no longer contained fat tails and was therefore closer to the measured distribution (compare Fig. S3c with Fig. 2b).

### *Ori* ensure accurate partitioning by perturbing the balance of fluxes

Let us review the factors determining positioning in the model (Fig. 4a). MukBEF self-organises via a Turing instability into regions of high local density (foci). The position of these foci is determined by the balancing of fluxes originating from a well-mixed (not self-organised) state. With spatially homogeneous conversion from this well-mixed state, the fluxes balance, in the case of a single focus, at the mid-cell position (i). An object (*ori*) that is attracted up the resulting gradient will also be positioned at mid-cell (Fig. 2). However, if the conversion from the well-mixed state is not homogeneous, then the position at which the fluxes balance changes (ii). If higher conversion (loading) occurs at the *ori*, then the mutual attraction between the self-organised MukBEF focus and the *ori* results in strong colocalisation (Fig. 3e) and positioning at mid-cell (iii). The system is analogous to a compound spring with the fixed end at mid-cell (but with a non-trivial relationship between the spring constants and specific loading ratio). At high levels of specific loading, colocalised MukBEF-*ori* becomes less sensitive to the (decreased) incoming flux from other regions, and is therefore less attracted towards mid-cell, resulting in more diffusive behaviour and a broader position distribution (Fig. 3f, S3a).

**Figure 4:**
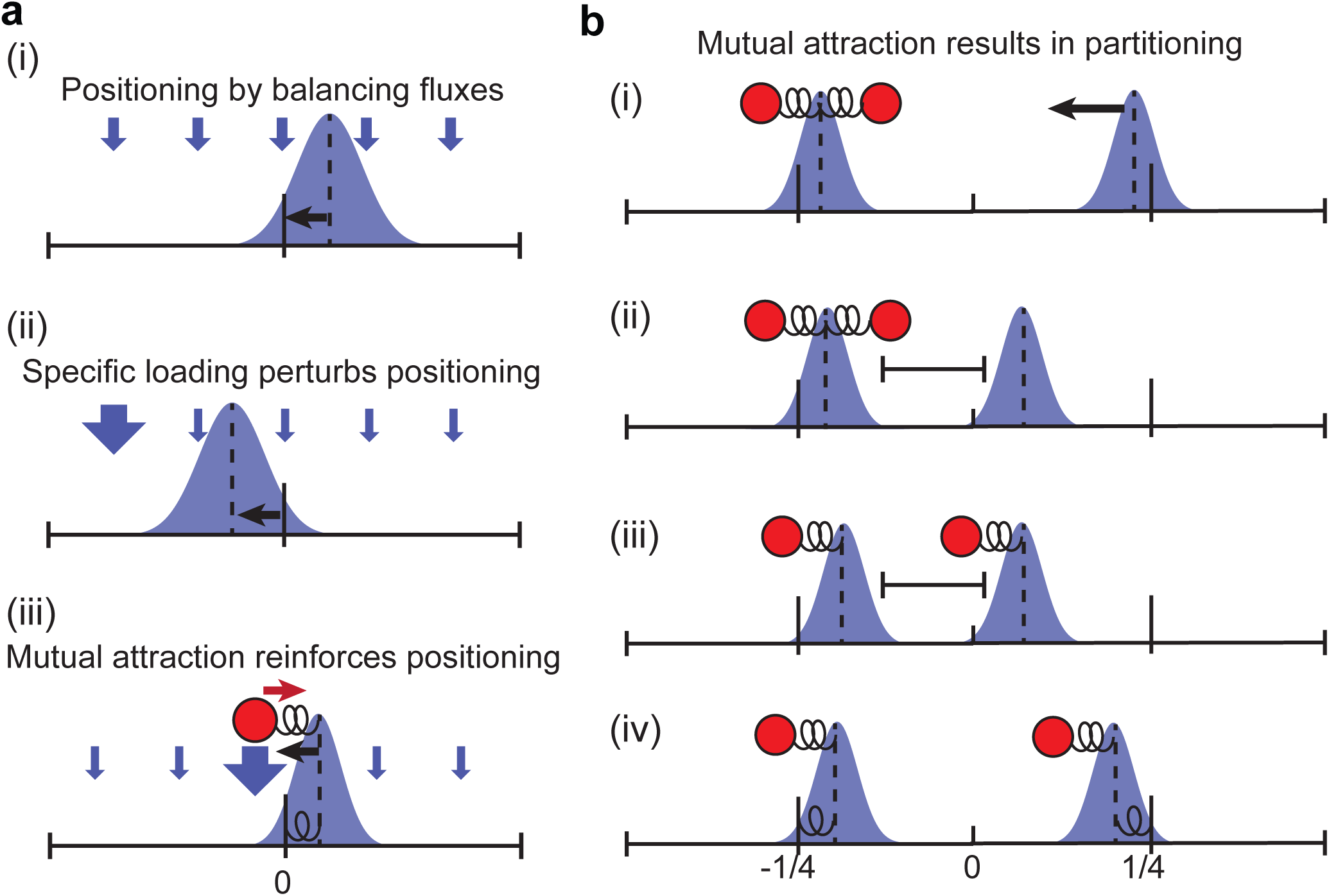
Schematic of positioning mechanism (a) With spatially homogeneous loading MukBEF peaks are positioned by regularly positioned by the balance of fluxes (i) [33]. Specific loading purtubs positioning (ii) and combined with the movement of ori up MukBEF gradient leads to robust positioning of both *ori* and MukBEF (iii). They remain in association, moving stochastically but in synch, behaving as if a single entity. (b) The undesirable configuration of both *ori* associated to the same MukBEF peak is unstable. The lone MukBEF peak is attracted towards the direction of higher flux (i-ii). Both peaks being closer, there is a higher chance that one of the *ori* jumps stochastically into the ‘basin of attraction’ of the lone peak (iii). The effect of balancing fluxes then causes each MukBEF-*ori* ‘complex’ to move toward their respective quarter positions (iv), the long-term stable configuration.

How do these properties lead to accurate and stable partitioning of *ori*? Suppose both *ori* have been pulled towards the same MukBEF focus (Fig. 4b). Due the flux imbalance, the MukBEF focus without an *ori* will be effectively pulled towards the other one (i) until the fluxes balance (ii). The stochastic nature of *ori* and MukBEF movement mean that at some point one *ori* leaves the influence of its MukBEF focus and becomes associated to the other focus (iii). When this occurs the balance of fluxes is again disturbed and MukBEF-*ori* move as single units towards opposite quarter positions (iv). This is the stable configuration.

### Timelapse microscopy supports the attraction of *ori* to MukBEF foci

The main experimental support that MukBEF positions *ori*, comes from the result that MukF depletion leads to *ori* mis-positioning which is restored upon repletion [30]. However, this could be an indirect effect of the resultant nucleoid decondensation. We therefore wanted to assess the relationship between MukBEF and *ori* without perturbing their positioning. If MukBEF and *ori* are positioned independently to mid-cell (in cells with a single *ori*) then we can predict the distribution of their separation from their individual distributions. This can then be compared to the corresponding measured distribution. To do this we first treated cells carrying fluorescently labelled MukB and *ori*, with DL serine hydroxamate so that they have a single non-replicating chromosome and therefore a single *ori*. Upon imaging, we found a substantial difference between the measured and predicted separation distributions (Fig. 5a,b). This indicates that MukBEF and *ori* are more colocalised then would be expected if they were positioned independently i.e. MukBEF and *ori* are not independently positioned. Furthermore, previous results indicate that MukBEF positioning is independent of *ori* positioning [10]. This suggests that either MukBEF positions *ori* or they are both positioned by the same unknown external mechanism.

**Figure 5:**
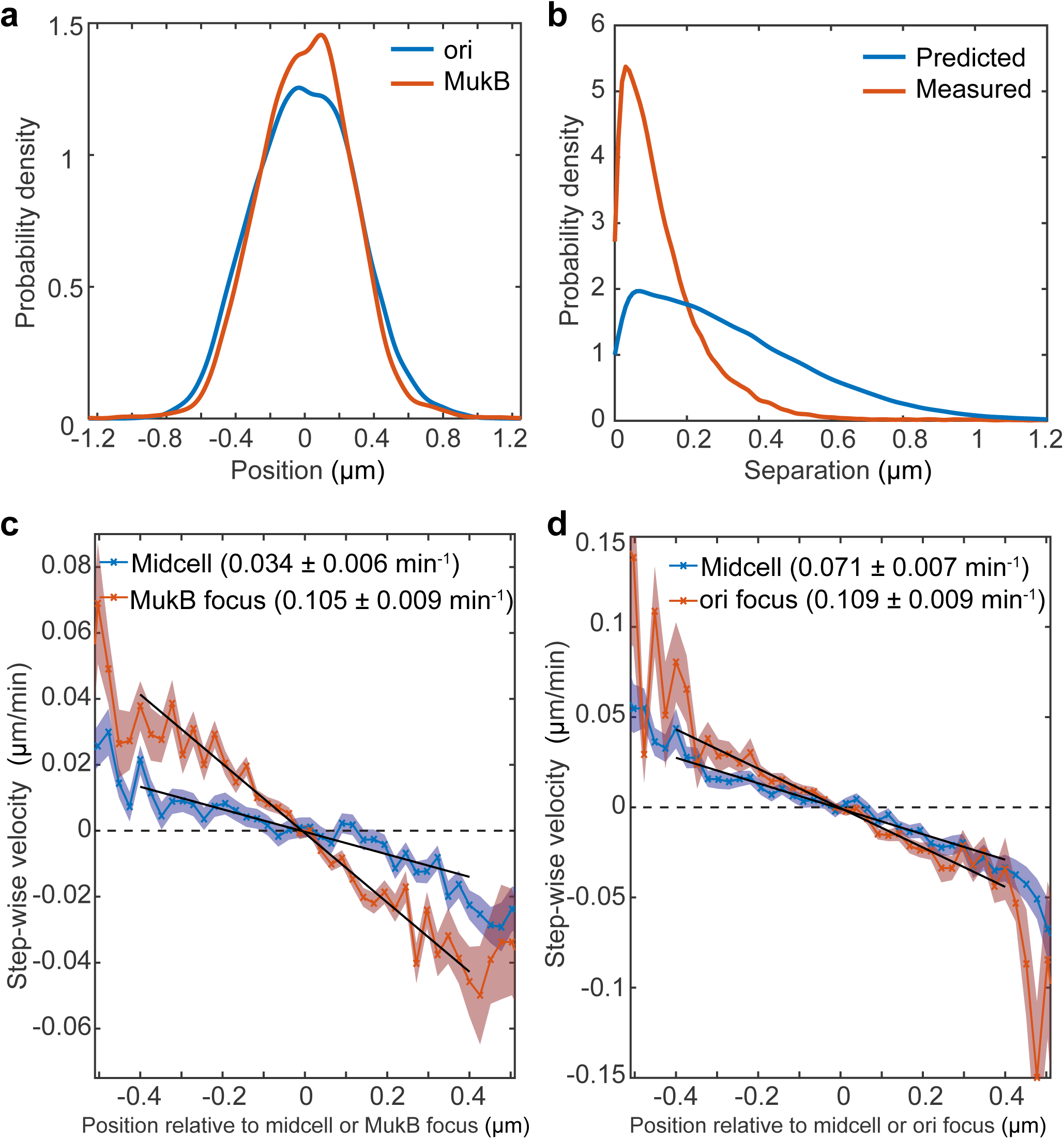
Fluorescence microscopy indicates that *ori* are attracted towards MukBEF. A strain with FROS labelled *ori* and MukB-mYPet was treated with DL serine hydroxamate to obtain cells with a single non-replicating chromosome and imaged at 1 min intervals. (a) The position distribution (along the long axis of the cell) of fluorescent foci of *ori* (blue) and MukB-mYPet (red). N=29374 from 866 cells tracked over up to 56 frames. (b) The predicted distribution (blue) of the distance between *ori* and MukB-YPet foci given that the distributions in (a) are independent. The measured distribution (red) of separation distances from the same cells. (c) and (d) The step-wise velocity of *ori* (c) and MukB-YPet (d) from the same cells as (a) and (b) measured as a function of position relative to midcell (blue) and the MukBEF (c) or *ori* (d) focus (red). Shaded regions indicate standard error. Black lines are weighted linear fits with the value of the slope and 95% confidence bounds indicated. Note that non-overlapping confidence intervals indicate the slopes are significantly different at the 5% level.

To distinguish between these two possibilities, we measured the step-wise velocity of *ori* as a function of its position relative to both mid-cell and the MukBEF focus. We found that *ori* shows a stronger restoring velocity towards to MukBEF than towards mid-cell (Fig. 5c). On the other hand, when we examined the dynamics of the MukBEF focus, we found a small but statistically significant bias towards *ori* (Fig. 5d). We found consistent results when we performed a similar analysis on the output of the simulations (Fig. S4). At low specific loading ratios, MukBEF is more strongly attracted to mid-cell (Fig. S4a,d), whereas at high ratios it is more strongly attracted to *ori* (Fig. S4c,f). At intermediate values the attraction is similar (Fig. S4b,e), consistent with the experimental observation (Fig. 5c,d). Together with the above results, these data suggests that *ori* movement is biased in the direction of the MukBEF focus, while the MukBEF focus is biased both towards mid-cell and *ori*, with the latter being potentially explained by the effect of specific loading within the *ori* region.

### Accurate partitioning during growth

We next incorporated exponential growth and stochastic *ori* replication into our simulations. We found that newly duplicated *ori* often remained associated to the same MukBEF focus, resulting in inaccurate partitioning as in the static case (Fig. S5). We also found that specific loading at *ori* delayed the splitting of MukBEF foci, similar to a spatially fixed loading site (Fig. S5b) [33]. However, in the subset of simulations in which partitioning occurred, *ori* were correctly positioned to the quarter positions.

We wondered how the system could be pushed out of the undesirable configuration with both *ori* associated to one MukBEF foci. One possibility has been suggested by polymer simulations. In the concentric shell model [42,43], nucleoid compaction facilitates the onset of segregation by promoting extrusion of newly synthesised DNA to the periphery of the nucleoid. The two *ori* experience the entropic repulsion of two closed loops and separating, initially rapidly, in opposite directions. This is consistent with more recent experimental observations. Duplicated *ori* (and indeed all loci) experience an initially large segregation velocity [14,44] followed by a slower phase in which they approach the quarter positions [13,14].

To mimic this repulsion of replicated *ori*, we incorporated a repulsive force into the simulations (see methods for details). We found that this was able to push the system sufficiently out of the undesirable configuration so that the system reaches the stable correctly partitioned configuration. The MukBEF focus splits, *ori* associate to separate foci and both move towards opposite quarter positions (Fig. 6a-d). The strength of the repulsion and the specific loading rate were chosen to give (at 140 min. simulation time) high (>95%) partitioning accuracy and an average *ori*-separation of half the cell length (see methods for details). Importantly, repulsion between *ori* without specific loading was not sufficient to ensure accurate positioning and partitioning (Fig. S6a,b).

**Figure 6:**
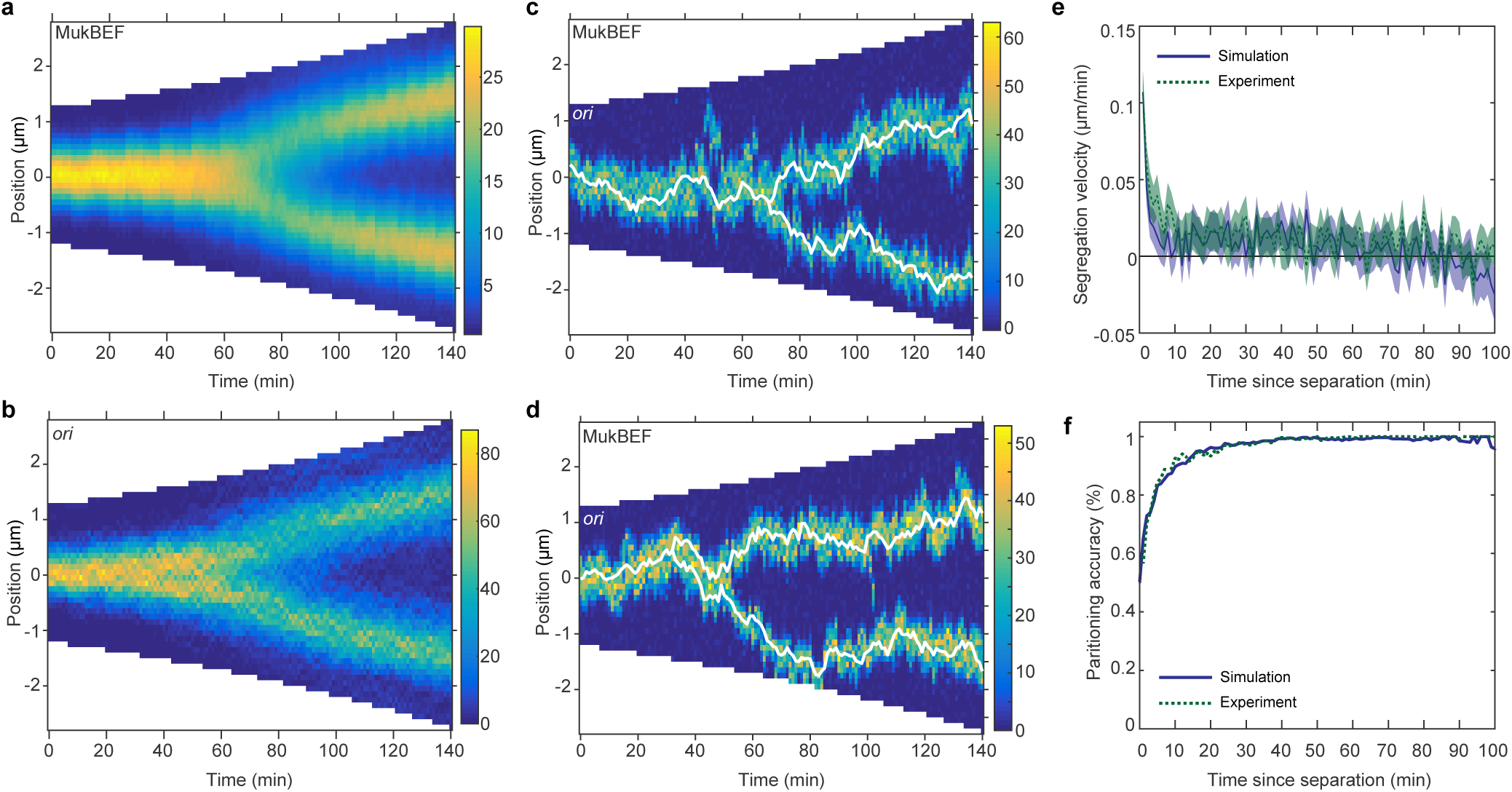
Repulsion between newly replicated *ori* promotes the transition to the stable configuration. (a) and (b) Average kymographs of MukBEF (a) and *ori* position (b) during exponential growth in the presence of a repulsive force between *ori*. (c) and (d) Two example kymographs from individual simulations (randomly selected) showing number of MukBEF molecules (colour scale) and *ori* position (while lines). (e) and (f) Segregation velocity (the step-wise rate of change of the absolute distance between *ori*) (e) and partitioning accuracy (f) plotted as function of the time since separation from simulations (blue) and experiment (green). Experimental data is from Kuwada et al. [13] and uses the synchronisation of that work. Shading indicates 95% confidence intervals. See methods for details of the synchronisation of simulated data. The segregation velocity has been corrected for growth. Simulation results are from 450 independent runs and used a doubling time of 120 min.

To compare the simulated time-courses to experiment, we synchronized the *ori* trajectories to the time of splitting as has been done for the experimental data [13]. We found excellent agreement of the segregation velocity (the change of the absolute distance between *ori* between time points) and the partitioning accuracy (Fig. 6e,f). We also measured *ori* separation as a function of time and found that it compared favorably but with some disparity, likely due to the broad distribution of cells lengths in the experiments compared to the fixed range used in the simulations (Fig. S6). Overall this good agreement, especially in the absence of any systematic fitting, suggests that *ori* positioning may indeed be explained by their interactions with a self-positioning MukBEF gradient.

## Discussion

The chromosomal origin of replication is dynamically positioned within many bacteria. While this can be explained in some bacteria by the presence of ParABS systems, in others, such as *E. coli*, the mechanisms are unknown. In this work, we presented a model for origin positioning in *E. coli* based on self-organising MukBEF, a prokaryotic SMC complex that has important roles in chromosome organisation andsegregation. MukBEF forms foci at the middle and quarter cell positions that show remarkable colocalisation with *ori* in wildtype cells [10,12]. Furthermore, while MukBEF foci are positioned independently of *ori* [10], they are required for correct *ori* positioning [30]. In previous work [33], we showed how the formation and positioning of MukBEF clusters can be explained by a Turing-type mechanism for self-organisation and positioning.

Here, we extended that model and showed that it could explain origin segregation and positioning and accurately reproduce detailed experimental results. The fundamental assumption of the model is based on the aforementioned relationship with *ori.* We assumed that, like SMC in *B. subtilis*, MukBEF is specifically loaded onto the DNA at specific sites within the *ori*. This could be the case if the *ori* (or proteins bound to it) has a higher affinity for MukBEF than other regions. The first consequence of this assumption is that the self-organised MukBEF gradient can act as a potential energy surface for *ori*, with the result that *ori* climb the MukBEF gradient in a directed manner. This is similar to the DNA relay model [34,45] of ParABS-based positioning and to Brownian ratchet models [46,47] in general. However, the situation here is simpler in that the protein gradient is already and spontaneously established and so *ori* positioning requires no further energy input, similar to the proposed bulk segregation of chromosomes by membrane-based protein gradients [48]. Furthermore, the mechanism was supported by polymer simulations (Fig. 1). It is also worth noting that MukBEF has been indicated [49–52] to have some ability to multimerize (beyond the dimer of dimer functional unit [32]). This could play a role in the interaction between MukBEF-associated *ori* and the broader MukBEF gradient. Lastly, we note the relationship between MukBEF and *ori* might be explained in other ways. In particular, *ori* could act as a ‘stop’ site for translocating MukBEF complexes. Indeed, initial simulations indicate this as a possibility, which we leave for future study.

This first gradient-climbing property alone was sufficient to explain the observed linear velocity profile in the one-*ori* case (Fig. 2) but was not able to reproduce accurate partitioning when two *ori* were present (Fig. 3a). However, we found that a higher relative loading rate of MukBEF within *ori* resolved this and achieved better agreement between the distributions of *ori* position (Fig. S3c,d) and *ori*-MukBEF separation distance (Fig. 3e) with the experimental results (Fig. 2). Since specific loading perturbs MukBEF foci positioning, pulling them towards the location of the loading site (Fig. 4a(ii)), it means that *ori* and MukBEF are effectively both attracted to each other (Fig. 4a(iii)). Each *ori*-MukBEF behaves as a single unit with *ori* essentially *‘*piggy-backing’ on MukBEF foci, using their intrinsic properties of positioning and mutual repulsion (Fig. 4b).

We next experimentally tested the model assumptions using wild-type cells. We found firstly that MukBEF and *ori* are not positioned independently of one another (Fig. 5a,b) and secondly, that *ori* display a significantly stronger restoring velocity towards MukBEF than towards midcell (Fig. 5c). On the other hand, and in agreement with the model, we found that MukBEF foci are similarly attracted towards both *ori* and midcell.

Finally, we assumed that newly duplicated *ori*s experience an effective repulsive force. This is not a new idea but is based on entropic repulsion between polymers [42,43,53]. In our model, this effect pushes the system out of a configuration with both *ori* associated to the same MukBEF clusters (the undesirable configuration), allowing it to fall into the desirable configuration having each *ori* associated to a different quarter-positioned MukBEF cluster. This configuration is stable and does not transition back to the undesirable configuration even in the absence of the repulsive force (Fig. 3). Furthermore, the repulsive force on its own is not sufficient for accurate segregation (Fig. S6). Specific loading at the *ori* is required as this is what leads to the stability of desired configuration (Fig. 4b). The repulsive force is necessary to get the system out of the undesirable state but not for the stability of the desirable state.

Comparing the model to experiment, we found excellent agreement. This is remarkable given the simplistic 1-D reaction-diffusion nature of the model and that we did not perform a systemic fitting of parameters to the experimental data (see methods for details of parameter selection). Furthermore the depth of the comparison is, to our knowledge, beyond what has been achieved previously for origin segregation in *E. coli*. Hence, we suggest that this approach warrants further consideration and that protein self-organisation may have an unappreciated role in chromosome organisation.

More generally, the idea of dynamically controlling the positioning and splitting of a Turing pattern is interesting from a mathematical point of view and may he applicable to other unrelated systems. Indeed, a major aspect of the ‘robustness problem’ of Turing patterning is the sensitivity of splitting to model parameters, domain size and stochastic effects [54]. The non-trivial coupling to *ori* in our model, as well as the self-positioning nature of the pattern [33], goes some way towards mitigating this sensitivity.

For future work, careful simultaneous measurement of *ori* and MukBEF dynamics would improve parameter estimation and provide ways to directly test the assumptions of the model. In the longer term, combining particle and polymer simulations may provide a deeper understanding of the process. In particular, MukBEF has a major role in chromosome organisation and facilitates long-range DNA interactions [40]. Like other SMC complexes, MukBEF may act by extruding loops of DNA [40], and/or be involved in stabilising them [52]. Furthermore, MatP, which binds to *matS* sites in the terminus region, interacts with MukBEF, displacing it from that region of the chromosome [12] and thereby restricting long-range DNA interactions between the terminus region and other regions of the chromosome [40]. Both effects may help position this region at mid-cell, while simultaneously encouraging the co-localisation of *ori* with MukBEF [12]. Thus, the role of MatP may need to be incorporated into future models. Overall, we envisage that the study of protein self-organisation in the context of chromosome dynamics has a promising future.

## Methods

### Review of the model

We briefly summarise the underlying model for MukBEF self-organisation [33]. The general model scheme consists of two ‘species’ *u* and *v* interacting in a bounded one-dimensional ‘bulk’ over a one dimensional surface (Fig. S1a, upper panel). Species *u* exists in the bulk (the cytosol) with concentration *u*_*c*_ and on the surface (the nucleoid) with concentration *u*_*n*_. Species *v* exists only on the surface with concentration *v*_*n*_. Species *v* diffuses slower than *u*. In the context of MukBEF, *u*_*n*_ represents the concentration of the free form of the basic functional complex, the dimer of dimers, while *v_n_* is the concentration of the slower diffusing form of the complex (due to entrapment of DNA). The former becomes the latter at a basal rate α and cooperatively in a cubic interaction at a rate β, and the latter becomes the former at a rate γ. Both of these species are nucleoid bound. The concentration of the cytosolic state is given by *u_c_*, which converts to and from the bound states with linear rates ⅅ and δ respectively. Figure S1a shows a diagram of the proposed model. See [33] for further details.

The two states *u_n_* and *v_n_* generate the Turing pattern and the kymographs and distributions shown in this work are of *v_n_* (which we simply refer to as MukBEF). The cytosolic state *u_c_* is responsible for the positioning of the pattern and is well-mixed because it interconverts with the Turing species on a sufficiently slow timescale. Note that in general this state is not required to be cytosolic, only well-mixed.

### Stochastic Simulations

Stochastic simulations are based on an exact realization of the Reaction Diffusion Master Equation (RDME) using the Gillespie method as described previously [33] but with changes for efficiency and the addition of simulated *ori*. We replaced the binary tree search of the enhanced direct method [55] with a 2D search as proposed by Mauch and Stalzer [56] and switched from 32-bit uniform random numbers (using the Ziggurat method) to 64-bit numbers (using std∷mt19937_64) to ensure enough significant digits to of the cell) is divided into discrete compartments, each having a width of *h*=0.1 μmand between which the species can diffuse. The accurately sample reactions occurring with very low relative rates (namely, *ori* diffusion). As before, the spatial domain (the long axis simulations were extended by the addition of *ori*, treated as an additional diffusing species, as follows.

#### Diffusion of *ori* subject to the MukBEF gradient

We modelled *ori* simply as diffusing particles. The drift they experience is based on the local MukBEF concentration and we use jump rates (the rate at which *ori* jump between neighbouring compartments) derived by Wang, Peskin and Elston [57]. The forward and backward jump rates from compartment to *i* to *i* + 1and *i*-1respectively are

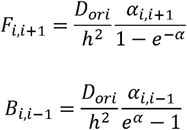

where 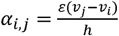 is, up to a factor, the difference in the MukBEF concentration between the compartments (*v_i_* is the number of molecules of slowly diffusing MukBEF species in compartment) *D* _ori_ is the diffusion constant and, is the drift parameter, determining the strength of the attraction up the MukBEF gradient. This form for the jump rates, which respects detailed balance, relies on the assumptions that, within individual compartments, the probability density for *ori* is at steady state and the MukBEF gradient is approximately linear. Both of these requirements can be satisfied for sufficiently small compartment widths. However, it is not feasible to decrease the compartment width much below 0.1 µm due to the increased computationally cost but also more fundamentally because of the intrinsic limitation of the RDME for non-linear reactions at short spatial scales [58,59]. Yet, the often sharp MukBEF profile (at a fixed moment in time) suggested that shorter compartment widths might be required. We therefore introduced sub-compartments within every compartment but only for *ori* positions. This approach has previously been applied to stochastic Turing patterns [60] but here we apply it to a ‘non-Turing’ species (*ori*). Each compartment was divided into an odd number of sub-compartments and the MukBEF concentration was linearly interpolated across sub-compartments. The jumps rates between sub-compartments were then defined as above. Performing simulations for different numbers of sub-compartments, we found that the apparent diffusion constant and drift rate (see below and Fig. 2) stabilised with greater than approximately 5 sub-compartments. The apparent diffusion was approximately 40% higher without sub-compartments. All results presented in this work used 21 sub-compartments as a higher value carried very little computationally cost. normal distribution with mean 3 μm and coefficient of variation 0.16 (based on the distribution of *ori*-foci splitting length of the data

#### Duplication of *ori*

The timing of *ori* replication was chosen randomly in each simulation by picking a duplication length from normal distribution with mean 3 μm and coefficient of variation 0.16 (based on the distribution of *ori-foci* splitting length of the data a in Kuwada et al. [13]). As we use a fixed growth range (2.5-5 µm) and doubling time (120 min), we truncate the distribution to this range. This duplication length is then converted to a duplication time via the exponential relationship between cell length and time. The simulation is paused when it reaches this time, the *ori* is duplicated (remaining within the same sub-compartment) and the simulation continued. Note that we do not mimic cohesion of newly replicated stands so that what we refer to in the simulation as *ori* duplication actually more closely corresponds to initial *ori* separation *in vivo*, which occurs 10-15 minutes after replication initiation.

#### Ori repulsion

As discussed in the text, newly duplicated *ori* are likely to experience, for entropic reasons, a repulsive force between them [42]. We implement this in the model by in the introduction of inverse square force between *ori*. This form of the force was chosen arbitrarily. However, we do not expect the precise nature of the force to affect our results as this force is only required to destabilise the undesirable configuration having both duplicated *ori* associated to the same MukBEF focus. It is not required for the _d_2are in the overdamped regime such that the separation velocity due to this force is proportional to the force i.e. 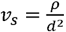, where *d* is the distance between *ori*. On the lattice, *d* is replaced by *d* _+_ *d* _o_, where *d* _o_ is the average distance between two uniformly distributed particles within a sub-compartment in order to the avoid a divergence when both *ori* are in the same sub-compartment. Dividing *v_s_*by the width of a sub-compartment (*h*_s =_*h*/21) defines the propensity function for the reaction that one of the *ori* (randomly chosen) moves one sub-compartment further away from the other *ori*.

#### Initialisation of simulations

Initial concentrations were set to the homogenous configuration (to the nearest integer). Unless stated otherwise, single *ori* were initially placed at mid-cell, while in simulations starting with two *ori*, they were placed at the quarter positions. Simulations were first run for 30 min to equilibrate and then read out every 1 min (chosen to match the experimental data). For simulations with growth, the simulation was paused after every time-duration that corresponded to growth by one compartment. An additional (empty) compartment was then inserted at a random position and the volume and total number of molecules (via the cytosolic fraction) were increased, maintaining the same overall concentration.

### Parameters

All parameters of the core MukBEF model are as previously described, except for the total species concentration, which is increased by 30%. This was done for compatibility of the MukBEF splitting time with the lower range of cell lengths used in this work (2.5 µm – 5 µm), which were chosen to more closely match the range of the experimental data in Kuwada et al. [13]. The remaining (*ori*-related) parameters were chosen by comparison with experimental data as described below.

### Apparent *ori* diffusion constant and drift rate

To be able to quantitatively compare the experimental and simulated data, we needed quantitative descriptors of the *ori* dynamics. We compared both data sets to a theoretical model of particle diffusion in a harmonic potential 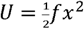 over an infinite 1D domain. Given a particle at position *x*_o_, the probability density that it is at position *x* at time δ*t* later is [61]

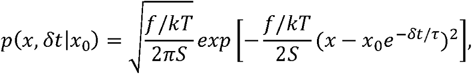

where s=1-e^-2δt/τ^ and 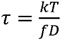 From this, it is straightforward to calculate the expected value and variance of the step-wise velocity, 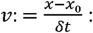:

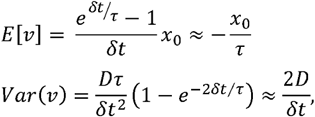

where the second equality holds for 2δt/τ≪1 (the full expression is used when fitting). Note the expected value of the step-wise velocity depends linearly on position, while the variance is independent of position. This is observed in both experiments and simulations close within the neighbourhood of the *ori* ‘home’ position. We therefore use the measured slope of the velocity relationship and its variance to determine an apparent diffusion constant, *D* _a_ and drift rate 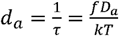

### Parameter fitting of *ori* related parameters

To search for parameter values for the *ori* diffusion constant (*D*_*ori*_) and the strength of attraction towards MukBEF (ε) that gave agreement between the measured apparent diffusion constants (*D*_*a*_) and drift rates (*d*_*a*_), we performed simple parameters sweeps. For the initial fitting (Fig. 2), we chose *D*_*ori*_ to be a percentage of the experimentally measured diffusion constant *D*_*a*_ ranging from 70% to 110% in 5% intervals, while the drift parameter ε was ranged from 0.5 to 2.5 times a nominal value of 0.026 µm (in steps of 0.5). Simulations were performed as described in Figure 2. The combination giving the best agreement and shown in Figure 2 was *D*_*ori*_ =0.9 *D* _a_ =5.4 x10^-5^µm2s-1 and, ε =0.026 µm. These values were also used for the simulations shown in Figures 3, S1, S2 and S3a,b.

To produce Figure S3c,d, we performed the same parameter sweep in the presence of 6x specific loading and found that the best agreement was obtained with, *D*_*ori*_ = 0.85, *D*_*ori*_ =5.1 x10^-5^µm^2^s^-1^ and ε =2xc0.026 µm (shown in Fig. S3c,d). It should be noted that given the stochastic nature of the simulations, even with 100 runs of 600 min each, there was quite some variability in the data. Furthermore, there appeared to a one parameter family of choices that give acceptable agreement in the diffusion and drift rates. However, they could be distinguished by the kurtosis (‘tailedness’) of the distribution of ori positions and we took this into account. These parameters were used for subsequent simulations (Fig. 6, S5,S6 and S7).

To find the optimal values for the specific loading ratio and the ori-repulsion rate constant (Fig. 6), we used a similar approach. We swept over specific binding ratios of b = [1,5,10,15,20] and repulsion rate constants of 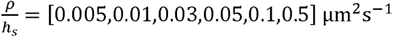. Simulations were performed as described in Figure 6. We examined the fraction of simulations that were correctly positioned and the average ori. We found that the combination of b=10 and 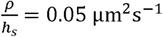 was the lowest valued combination that displayed greater than 95% partitioning accuracy and a separation of approximately 50% of the cell length.

To examine the dynamics of new replicated *ori*, we synchronised the simulations to the time of *ori* duplication. We measured the segregation velocity of *ori* (the step-wise rate of change of the absolute distance between *ori*) across the synchronised simulations and found very good agreement with a similar analysis of the experimental data (Fig. 6e). The parameters of the model and the *ori*-duplication length have not been chosen such that, for a given specific loading ratio, the duplication of *ori* is coincident with the length-dependent splitting of the MukBEF focus (due to the computational complexity of such a task). Indeed, *ori* duplication occurs on average at approximately 37 min into the simulation, notable earlier than MukBEF splitting (Fig. 6). Thus, for longer timescale *ori* dynamics, we synchronise the simulations to the last time point at which the *ori* are within 500 nm of each other (results are not very sensitive to the precise value). When we did this we found excellent agreement in the partitioning accuracy and *ori* separation between simulations and experiment (Fig. 5f, S7). With synchronization to *ori* duplication, the *ori* separation profile was sigmoidal with an initial plateau due to both *ori* being in close association with the same and usually only MukBEF focus (as in Figure 6d). Thus, synchronizing as we have done ensures that we assess the dynamics from the time of separation rather than the time of duplication, which is in keeping with the experiments.

Analysis, fitting and plotting were performed in MATLAB (Mathworks Inc.)

### Polymer simulations

In order to investigate the interplay between MukBEF and *ori* positioning within the nucleoid, we used a coarse-grained lattice polymer [41]. Within this framework, the polymer is described as a self-avoiding ring polymer that is confined in an elongated cuboid with an aspect ratio of 4:1 comparable to that of the *E. coli* nucleoid. Using a ring polymer composed of 464 monomers, we chose a lattice of size 56 x 14 x 14 that leads to a system density (monomer-to-volume ratio) of around 10%. Dynamic looping interactions were enabled between one specific monomer (*ori*) and distant monomers. The looping probability is set to be dependent on the spatial position of *ori* along the long axis of the nucleoid and is drawn from a Gaussian distribution centred around mid-cell with a standard deviation of 5.6 lattice units. A lifetime of 10000 Monte-Carlo steps (MCS) was assigned to each loop. 10 independent Monte-Carlo trajectories were used to sample the dynamics of the system. In each simulation, 10000 polymer conformations were recorded, one every 50000 MCS. The initial position of *ori* was varied in each simulation in order to uniformly cover the long axis of the cuboid.

### Experiments

Strain SN192 (AB1157 *lacO240∷hyg* at *ori1, tetO240∷gen* at *ter3, Plac-lacI-mCherry frt* at *leuB, Plac-tetR-mCerulean frt* at *galK, mukB-mYPet frt*) [12] was grown in M9 minimal medium supplemented with 0.2% glycerol and required amino acids (threonine, leucine, proline, histidine and arginine—0.1 mg ml^-1^) at 30 °C. Cells were grown O/N, diluted 1000-fold and grown to an A_600_ of 0.05–0.2. Unlike longer cells with two (quarter positioned) *ori* foci, cells with a single (mid-cell localised) *ori* focus, can be analysed together in absolute, rather than scaled, coordinates by simply measuring foci positions relative to mid-cell (as we did in Fig. 2 for the dataset of Kuwada et al. [13]). We therefore treated cells with DL serine hydroxamate (Sigma-Aldrich, S4503) to a final concentration of 1 mg/ml. During the treatment, cells do not initiate a new round of replication, but complete any ongoing rounds [62]. To allow sufficient time for replication to complete to termination, cultures were grown for an additional 3 h (generation time ∼170 min). Finally, cells were spotted onto an M9-gly 1% agarose pad on a slide for imaging.

Time-lapse movies were acquired on a Nikon Ti-E inverted fluorescence microscope equipped with a perfect focus system, a 100× NA 1.4 oil immersion objective, an sCMOS camera (Hamamatsu Flash 4), a motorized stage, and a 30 °C temperature chamber (Okolabs). Fluorescence images were automatically collected at 1-min intervals for 56 minutes using NIS-Elements software (Nikon) and an LED excitation source (Lumencor SpectraX). Exposure times were 150 ms for mCherry, and 100 ms for mYPet using 50% LED intensity. Phase contrast images were also collected at 1-min intervals for cell segmentation.

Alignment of frames, cell segmentation and linking of cells in consecutive frames were performed using SuperSegger [63]. To ensure that we considered only cells with a single *ori*, we subsequently filtered the dataset as follows. For any cells that had two *ori* foci on any frame, we kept only the frames before two foci were first detected. This reduced the dataset from 1311 to 866 cells. We then used only frames with exactly one *ori* focus and one MukB focus. This resulted in 29374 data points (cell-frame combinations). The distribution in Fig. 5a,b were made using the smoothening kernel estimate function *ksdensity* in MATLAB. Similar results were obtained if we further restricted the data set to cells of similar length. The same data was used to generate the step-wise velocity profiles (Fig. 5c,d) but as two consecutive frames are required, this reduced the data set to 24267 data points. Linear fitting (Figs. 2c, 5c,d, S3) was performed using the *fit* function with the inverse square of the standard errors as weights.

## Acknowledgments

We thank Remy Colin for discussions and Victor Sourjik for discussion, support and comments on the manuscript. This work was supported by the German Federal Ministry of Education and Research and the Max Planck Society in the framework of the MaxSynBio research network. Work in the D. J. Sherratt laboratory was supported by a Wellcome Investigator Award (DJS: 200782/Z/16/Z).

**Figure Legends**

**Figure S1** Properties of MukBEF model and *ori* positioning. (a) Schematic showing the reactions of the system (see methods). Species *u* exists in the bulk (red) or on the surface (green). Species *v* exists only on the surface (blue). Binding and species interaction are indicated by arrows. Diffusion is not shown. (b) The average MukBEF profile in the simulations (blue dots) is well approximated by a Gaussian. (c)Average kymographs from 100 simulations showing the distributions of MukBEF and *ori* positions. (d) Cumulative probability distribution as in Fig. 3e but for simulation with one *ori*. Simulations were performed as in Fig. 2.

**Figure S2** Additional properties of model in longer cells with two *ori.* (a) Histograms of MukBEF numbers for 1x, 6x and 16x specific loading. Positioning is more precise at 6x than with no or 16x specific loading. (b) Fraction of simulation time with two MukBEF foci as a function of the specific loading ratio. Error bars indicate standard deviation. (c) Partitioning accuracy and stability as in Fig. 3c but using random initial *ori* positions. Data is from the last 100 min of 600 min simulations. Due to both *ori* sometimes being in close initial proximity, the occurrence of undesirable configuration (both *ori* associated to the same MukBEF peak) is higher. Nevertheless the system eventually tends to transition (irreversibly) into the correct configuration, promoted by high specific loading rates. The loading ratios are as in (b).

**Figure S3** Additional properties and re-parameterisation of model in short cells with one *ori.* (a) The variance of the MukBEF and *ori* position distributions for different specific loading ratios for the case of a single *ori*. Simulations as in Fig. 2. (b) The change in the apparent diffusion constant and drift rate as a function of the specific loading rate. (c) and (d) A refitting of the simulated data in the presence of 6x specific loading. Otherwise as in Fig. 2b,c. Compare with experimental result in Fig. 2b,c. See methods for further details and model parameters.

**Figure S4** MukBEF is attracted towards midcell at low loading ratios and towards *ori* at high loading ratios. (a-c) Simulated position distributions of MukBEF and *ori* for no (a), six times (b) and 12 (c) times specific loading. The MukBEF position is the centroid of the simulated MukBEF peak. (d-f) Simulated step-wise *ori* (top) and MukBEF velocities (bottom) for no (d), six times (e) and 12 (f) times specific loading as above. Otherwise as experimental data in Fig. 5c,d. Shaded region indicates standard error. Compare (b) and (e) with Fig. 5a,c,d. Note the inversion in the dominant MukBEF restoring velocity as specific loading is increased. The position distribution and *ori* step-wise velocity for the case of no specific loading are reproduced from Fig. 2b,c. Note that MukBEF is not affected by *ori* in the absence of specific loading. The apparent restoring velocity of MukBEF towards *ori* in the bottom panel of (d) is not physical and is likely due to the higher variability (diffusivity) of the MukBEF focus compared to *ori*.

**Figure S5** The cycle-cycle dependent effect of specific loading. (a) and (b) Average kymograph of the number of MukBEF molecules and *ori* positions during exponential group with homogeneous loading (a) and 10x specific loading at *ori* (b). Partitioning accuracy is measured by the proportion of simulations having partitioned *ori* at the end of the simulation (140 min). (c) and (d) Two randomly selected individual simulations from (a) and (b) respectively. The colour scale represents the number of MukBEF molecules; white lines show the positions of *ori*. Simulation results are from 450 independent runs and used a doubling time of 120 min.

**Figure S6** Repulsion between *ori* is not sufficient for correct segregation and positioning. (a) Average kymograph of the number of MukBEF molecules and *ori* positions during exponential group with no specific loading but with repulsion between *ori*. Partitioning accuracy is measured by the proportion of simulations having partitioned *ori* at the end of the simulation (140 min). (b) Two representative individual simulations from (a). The colour scale represents the number of MukBEF molecules; white lines show the positions of *ori*. Simulation results are from 450 independent runs and used a doubling time of 120 min. See Fig. 6 for the effect of repulsion in the presence of specific loading. The strength of the repulsive force is the same as in Fig. 6.

**Figure S7** The *ori* separation distance is in qualitative agreement with experimental results. The *ori* separation distance relative to cell length plotted as function of the time since separation. Experimental curve obtained using the data of Kuwada et al. [13]. Simulated data is as shown in Fig. 6. Individual simulations were synchronised to the last time point for which the *ori* are within 0.5 µm of each other (as for partitioning accuracy in Fig. 6f). See methods for details. Shading indicates 95% confidence intervals.

